# Identification of Novel FosX Family Determinants from Diverse Environmental Samples

**DOI:** 10.1101/2024.11.04.621841

**Authors:** Nicolas Kieffer, Maria-Elisabeth Böhm, Fanny Berglund, Nachiket P. Marathe, Michael R Gillings, D. G. Joakim Larsson

**Affiliations:** Molecular Basis of Adaptation Laboratory, Departamento de Sanidad Animal, Facultad de Veterinaria de la Universidad Complutense de Madrid, Madrid, España; Centre for Antibiotic Resistance Research (CARe)in Gothenburg, Sweden; Department of Infectious Diseases, Institute of Biomedicine, Sahlgrenska Academy, University of Gothenburg, Gothenburg, Sweden; Department of Contaminants and Biohazards, Institute of Marine Research (IMR), Bergen, Norway; ARC Centre of Excellence in Synthetic Biology, Macquarie University, Sydney, NSW 2109, Australia; Department of Molecular Sciences, Macquarie University, Sydney, NSW 2109, Australia

**Author notes:** Corresponding Author., Department of Infectious Diseases, Institute of Biomedicine, The Sahlgrenska Academy at the University of Gothenburg, Guldhedsgatan 10, SE-413 46, Göteborg, Sweden.

## Abstract

**Objectives:** This study aimed to identify novel fosfomycin resistance genes across diverse environmental samples, ranging in levels of anthropogenic pollution. We focused on fosfomycin resistance, and given its increasing clinical importance, explored the prevalence of these genes within different environmental contexts.

**Methods:** Metagenomic DNA was extracted from wastewater and sediment samples collected from sites in India, Sweden, and Antarctica. Class 1 integron gene cassette libraries were prepared, and resistant clones were selected on fosfomycin-supplemented media. Long-read sequencing was performed, followed by bioinformatics analysis to identify novel fosfomycin resistance genes. The genes were cloned and functionally characterized in *E. coli*, and the impact of phosphonoformate on the enzymes was assessed.

**Results:** Four novel fosfomycin resistance genes were identified. Phylogenetic analysis placed these genes within the FosX family, a group of metalloenzymes that hydrolyse fosfomycin without thiol conjugation. The genes were subsequently renamed *fosE2, fosI2, fosI3*, and *fosP*. Functional assays confirmed that these genes conferred resistance to fosfomycin in *E. coli*, with with MIC ranging from 32 μg/ml to 256μg/ml. Unlike FosA/B enzymes, these FosX-like proteins were resistant to phosphonoformate inhibitory action. A *fosI3* homolog was identified in *Pseudomonas aeruginosa*, highlighting potential clinical relevance.

**Conclusions:** This study expands the understanding of fosfomycin resistance by identifying new FosX family members across diverse environments. The lack of phosphonoformate inhibition underscores the clinical importance of these poorly studied enzymes, which warrant further investigation, particularly in pathogenic contexts.

## 1. Introduction

Fosfomycin, an antibiotic discovered in the 1960s, is increasingly used to treat infections caused by multidrug-resistant bacteria. It is particularly effective in combination with other antimicrobials, due to its broad-spectrum efficacy and excellent bioavailability when administered intravenously. Fosfomycin is active against many Gram-negative and Gram-positive bacteria, and acts by inhibiting MurA (N-acetylglucosamine enolpyruvyl transferase), the enzyme responsible for the initial step in peptidoglycan synthesis, thereby disrupting bacterial cell wall construction [1].

Resistance to fosfomycin arises through various mechanisms. One mechanism involves point mutations in key genes for membrane transporters such as *glpT, uhpT*, and *uhpA*. These transporters are responsible for the uptake of the antibiotic, and changes to their structure lead to increased minimum inhibitory concentrations (MICs). Mutations in those genes can confer resistance to fosfomycin, but they often come at a fitness cost to the bacteria. This is because these transporters are responsible for the uptake of important nutrients like glycerol-3-phosphate and glucose-6-phosphate. Loss of function weaken cells by depriving them of these essential compounds, making them less competitive in environments without fosfomycin pressure [2]. Bacteria can also acquire plasmid-encoded fosfomycin resistance genes whose products are enzymes that inactivate the antibiotic. Among these, the FosA enzyme, a member of the glutathione-S-transferase (GST) family, is predominantly found in Gram-negative bacteria, while the FosB, part of the bacillithiol-S-transferase (BST) family, is common in Gram-positive bacteria. These enzymes catalyze the conjugation of a thiol group to fosfomycin, effectively neutralizing its antimicrobial activity [2].

In addition to these thiol-based resistance mechanisms, another group of mobilized genes known to encode FosX-like enzymes also contributes to fosfomycin resistance. Unlike the GST and BST families, FosX-like enzymes do not utilize a thiol group for inactivation. Instead, they function as metalloenzymes that directly hydrolyse fosfomycin, rendering it inactive. The *fosX* gene is the prototype of this family and is found across a diverse range of bacterial species, both Gram-positive and Gram-negative but was initially identified in the chromosome of *Listeria monocytogenes* [3].

Functional metagenomics is currently one of the most effective methods for detecting new resistance determinants before they become widespread in pathogens. This approach facilitates the discovery of new genes associated with known resistance mechanisms and enables the identification of entirely novel resistance mechanisms. In previous studies, we successfully identified new resistance genes within class 1 integrons using a PCR-based amplification method coupled with functional metagenomics [4–6]. The focus on class 1 integrons is explained by the fact that these genetic platforms are known to carry a myriad of gene cassettes, many of which contain antibiotic resistance genes, including those for fosfomycin [7–9].

Building upon our prior functional metagenomics studies, this study specifically focused on fosfomycin resistance, due to its crucial role as an antibiotic in contemporary medical treatments. Our investigation extends to various environmental contexts, from Antarctic environments to municipal wastewater from the inlet of the Rya municipal wastewater treatment plant of Gothenburg, Sweden, and highly polluted surface water from India. The aim of this study was to identify novel fosfomycin resistance determinants across different environmental settings and functionally characterize them to assess their potential clinical relevance.

## 2. Materials and Methods

### 2.1 Metagenomic DNA Samples

Samples were collected from four locations. Two polluted water samples were obtained from rivers in India: one from the Mutha River in Pune, Maharashtra (73°53’29.62’’ E, 18°32’35.05’’ N), and the other from the Nakka vagu River near the Patancheru Enviro Tech Ltd (PETL) wastewater treatment plant in Hyderabad (78°14’37.6’’ E, 17°32’23.1’’ N). Detailed descriptions of the Indian sampling process can be found in our previous study [10]. In addition, a wastewater sample was collected in December 2019 at the Rya wastewater treatment plant in Gothenburg, Sweden (11°53’31.2” E, 57°41’45.8” N). This sample consisted of a 24-hour composite of the plant’s influent. Lastly, soil samples were collected near the Casey Antarctic base (110°39’17.4’’ E, 66°24’36.8’’ S). For the water and wastewater samples, 500 ml of sample was filtered through 0.22 μm hydrophilic filters. In the case of the sediment sample from Antarctica, 0.3 g of soil was used directly for DNA extraction as described previously [11]. DNA was extracted from both the filtered water samples and the soil sample using the Qiagen DnEasy Powersoil kit. The quality of the extracted DNA was verified using Nanodrop and Qubit spectrophotometry.

### 2.2 Class 1 integron cassette library preparation

Class 1 integron gene cassette libraries were prepared and screened according to the protocol published previously [12] using the primers HS458 (5’-GTTTGATGTTATGGAGCAGCAACG-3’) and HS459 (5’-GCAAAAAGGCAGCAATTATGAGCC-3’). These primers amplify entire gene cassette arrays by binding to the 5′ and 3′ conserved segments of class 1 integrons. The vector pZE21-MCS1 (Expressys, Germany) was modified by inserting the constitutively active promoter P_*bla*_ and its ribosomal binding site via the restriction sites *KpnI* and *HincII*. The modified plasmid, pZE21-P_*bla*_, was then linearized using Phusion High-Fidelity DNA Polymerase (Thermo Fisher Scientific, USA) with the primers pZE21-for (5’-GACGGTATCGATAAGCTTGAT-3’) and pZE21-Pbla-rev (5’-GACTCTTCCTTTTTCAATATTATTGAA-3’) and subsequently dephosphorylated using FastAP (Thermo Fisher Scientific, USA). This approach allowed efficient library preparation and screening for mobilized novel resistance determinants. DNA from the samples was amplified using 5’ phosphorylated primers targeting the gene cassette array of class 1 integrons to favour ligation, as the plasmid vector was dephosphorylated.

Libraries of class 1 integron gene cassettes were constructed with overnight ligation using the Fast Link ligation kit (Epicentre, Lucigen, USA). Linearized pZE21-P_bla_ plasmids were ligated with gene cassettes amplified from the metagenomic DNA samples at a 1:5 molar ratio, followed by electroporation into *E. coli* DH10β. After recovery, aliquots of 1 µl, 0.1 µl, and 0.01 µl from each library were plated on LB agar with 50 µg/ml kanamycin to determine library size, plate count, and average insert size. The remaining libraries were inoculated into 10 ml LB with kanamycin and incubated at 37 °C, 180 rpm for 18 hours. Libraries were then aliquoted and stored in 20% glycerol at -80 °C. Library sizes were calculated by multiplying the average insert size by the number of colony-forming units after transformation recovery. Insert size distribution was assessed by PCR amplification and gel electrophoresis of inserts from 10 randomly selected clones from each library. For functional selection, 100 µl of each metagenomic library was plated on Luria Bertani plate agar supplemented with kanamycin (25 μg/ml) and with fosfomycin at 4x, 8x minimal inhibitory concentration (MIC), and/or the clinical breakpoint concentration for *E. coli* DH10β. This resulted in using plates supplemented with 4 μg/ml, 8 μg/ml, or 32 μg/ml of fosfomycin, respectively. Plates were incubated at 37 °C for 16-24 hours.

All resistant colonies from a single plate were harvested individually using a sterile disposable cell scraper (Sarstedt, Germany), pooled, and then resuspended in 500 μl of LB + 20 % glycerol and frozen at -80°C.

### 2.3 Amplicon-PCR and sequencing

The selected resistant clone libraries were thawed and 300 μl were washed twice in PBS buffer. Cells were subsequently pelleted a third time in nuclease-free H_2_O, re-suspended in 50 µl H_2_O and used as PCR template. For plates with 1-100 colonies, 1-2 μl of plate scrape lysate was used; for plates with 100-2000 colonies, 0.1 μl was used as template. Amplicons were prepared with Phusion High-Fidelity DNA Polymerase (Thermo Fisher, Gothenburg, Sweden) and primers binding directly up- and downstream of the insertion site within pZE21-P_*bla*_. The amplicon PCR primers contained sample specific barcodes that allowed allocation to the antibiotic and concentration used for selecting clones. The resulting PCR products were purified, quantified using Qubit® Fluorometer and quality was assured by Nanodrop^™^ spectrophotometer. Sequencing libraries were prepared from each pool using SMRTbell^™^ Template Prep Kit 1.0-SPv3 and the two libraries were sequenced on separate PacBio Sequel^™^ SMRT^®^ cells in the Science for Life Laboratories (Uppsala, Sweden).

### 2.4 Bioinformatics analysis

The raw sequencing data obtained from the PacBio Sequel™ system were first subjected to quality control using the PacBio SMRT Analysis software, which included filtering subreads to remove low-quality sequences and trimming adapters using pbccs v4.02 (https://github.com/PacificBiosciences/ccs) and BAM2fastx tool (https://github.com/pacificbiosciences/bam2fastx/). High-quality PacBio reads were imported into the SMRT Analysis software for initial quality control, including filtering subreads and adapter trimming. The cleaned reads were assembled into contigs using the Canu assembler [13], specifically designed for long-read sequences, followed by error correction with the Arrow tool [14]. The polished contigs were uploaded to the CARD database [15] via the RGI tool to identify and annotate resistance genes. These annotations were cross verified using ResFinder to ensure accuracy and provide additional context [16].

The identified *fos* homologs were extracted for further analysis, including comparison with known sequences in the CARD and ResFinder databases to assess novelty and functional implications. Phylogenetic analysis was conducted to determine the evolutionary relationships of the newly identified *fos* genes with existing *fos* gene families and phylogenetic trees were performed using the Seaview software [17] after collecting all characterized *fos* genes including bacillithiol, gluthatione-S-transferases and fosfomycin hydrolases.

Three-dimensional structures of the newly identified Fos proteins were predicted using the AlphaFold2 pipeline [18], specifically implemented through ColabFold[19]. Subsequently, Foldseek [20] was employed for an additional layer of comparison, focusing on aligning the protein structures rather than just their amino acid sequences.

### 2.5 Functional verification of novel *fos* genes and antimicrobial susceptibility testing

New *fos* candidates were synthesized in the pMK plasmid by Gene Art Gene Synthesis service (Thermo Fisher Scientific, Germany) as described previously [6]. Each gene sequence was flanked by the P_*bla*_ constitutively active promoter plus ribosomal binding site upstream and the *rrnB* terminator T1 downstream, respectively. The recombinant plasmids were electroporated into *E. coli* DH10β along with the empty pMK plasmid as a negative control.

Fosfomycin MIC was determined by agar dilution as recommended by the EUCAST guidelines [21]. Glucose-6-phosphate was added at a concentration of 25 μg/ml to Muller-Hinton agar plate with fosfomycin concentration ranging from 2 to 2,048 μg/ml. Phosphonoformate was added at a concentration of 5 mM to assess its inhibitory effect on *fos* genes as described previously [22].

## 3. Results and discussion

### 3.1 Detection of new *fos* determinants

The ligation of metagenomic DNA from all four sampling sites resulted in the emergence of fosfomycin-resistant colonies on LB plates supplemented with 8 μg/ml and 32 μg/ml of fosfomycin. The number of colonies obtained was 101 and 21 for Casey, 50 and 7 for Pune, 132 and 54 for PETL, and 150 and 35 for Gothenburg. Following amplicon PCR and sequencing, bioinformatics analysis identified a unique *fos* gene variant in each sample. BLAST analysis indicated that the encoded Fos proteins shared significant amino acid identity, ranging between 71.97% and 81.95% (Figure 1A), although these proteins had not been characterized previously.

**Figure 1.**
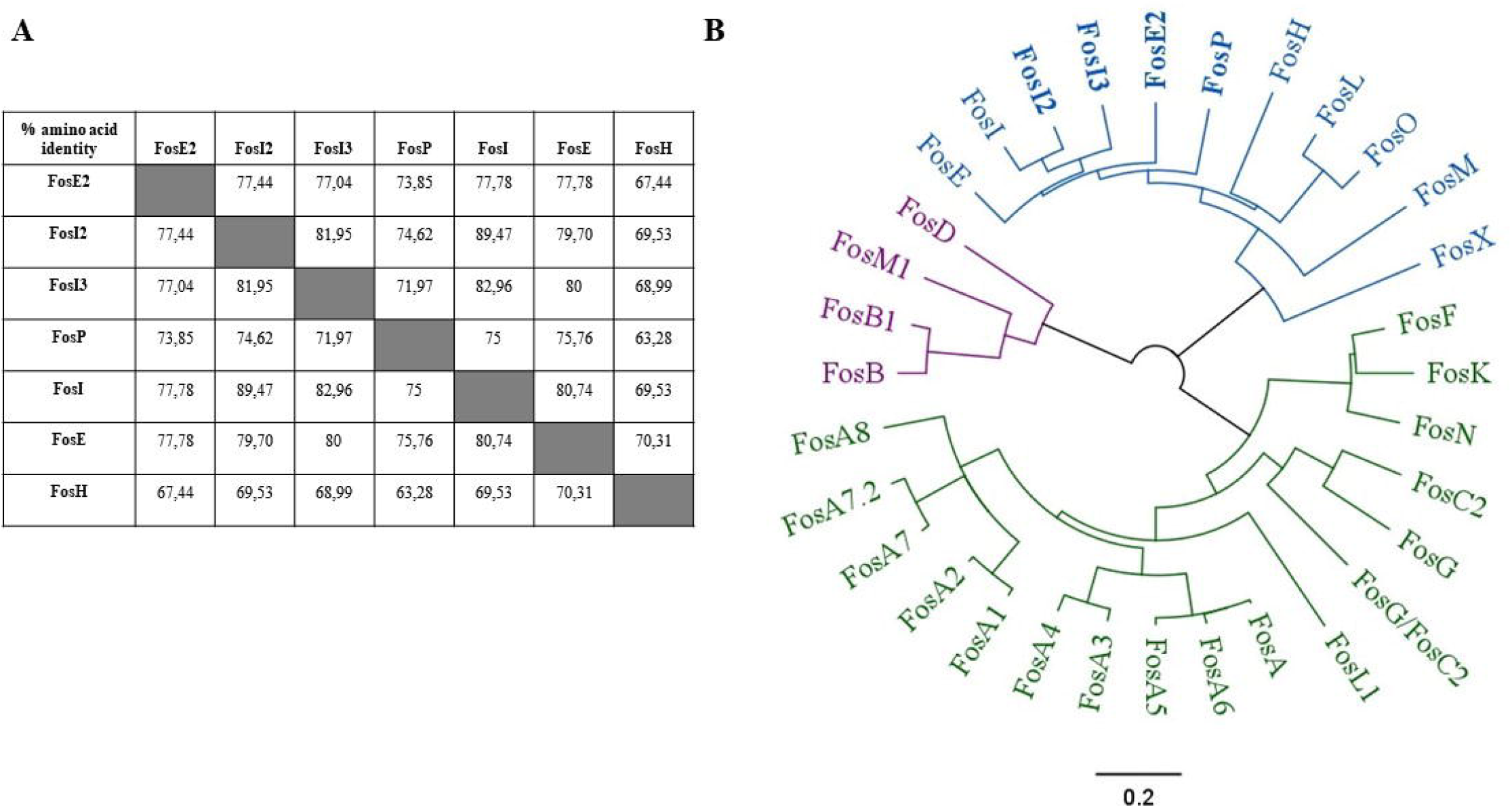
(A) Sequence identity matrix showing the percentage of amino acid identity between the different Fos proteins. Higher identity percentages indicate closer evolutionary relationships. Notably, FosI2 and FosI3 share high identity with FosI, reflecting their close relationship within the FosI subfamily. The same principle was applied to the identification of FosE2. (B) Phylogenetic tree illustrating the evolutionary relationships among the Fos proteins. The clustering of FosP in a distinct branch supports its classification as a new variant within the FosX family, with notable divergence from FosI and FosE. The blue color indicates enzymes belonging to fosfomycin hydrolases, while purple and green colors represent Fos proteins belonging to bacillithiol and glutathione transferases, respectively.

Further analysis was conducted using all characterized sequences of *fos* genes available in the NCBI database. Phylogenetic analysis demonstrated that the four new *fos* variants belonged to the same subtree as FosI and FosE (Figure 1B). Notably, the two Fos proteins recovered from Indian samples showed high identity with FosI (89.47% and 82.96%, respectively), indicating a close evolutionary relationship. Consequently, these proteins were named FosI2 and FosI3, respectively.

The Fos enzyme identified from the Antarctic samples, showed significant identity with FosE (77.78%), and displayed enough distinctiveness to warrant a new identifier. Thus, it was renamed FosE2, following the convention of using a numerical suffix for closely related but distinct variants. Finally, the protein encoded by the *fos* gene identified in Gothenburg clustered within the fosfomycin hydrolase subtree, but exhibited notable divergence from other members, particularly in evolutionary distances observed (Figure 1B). Given its moderate identity to both FosI (75%) and FosE (75.76%), and its distinct branch on the phylogenetic tree, we decided to assign a new designation, *fosP*, following the alphabetical convention commonly used for naming *fos* genes. AlphaFold analysis corroborated by FoldSeek alignment revealed that all newly characterized Fos enzymes aligned well with the FosX enzyme family, exhibiting Template Modeling Scores (TM-score) greater than 0.97 and Root Mean Square Deviations (RMSD) less than 0.8 Å, except for FosP which displayed an RMSD of 0.85 Å, as detailed in Supplementary Figure S1. This high level of structural similarity confirms these enzymes belong to the FosX family. This enzyme family includes FosE and FosI-like proteins, and are enzymes with functions akin to the metalloenzymes that hydrolyse fosfomycin without involving thiol conjugation [3]. This hydrolytic action is mediated by metal ions such as manganese, which act as crucial cofactors. Thus, while all these enzymes share a dependency on metal ions, the FosX family uniquely catalyses the breakdown of the antibiotic molecule itself, rather than modifying it through thiolation. Sequence alignment shows that all Fos enzymes described in this study shared the amino acid responsible for the hydrolytic activity (Figure 2).

**Figure 2.**
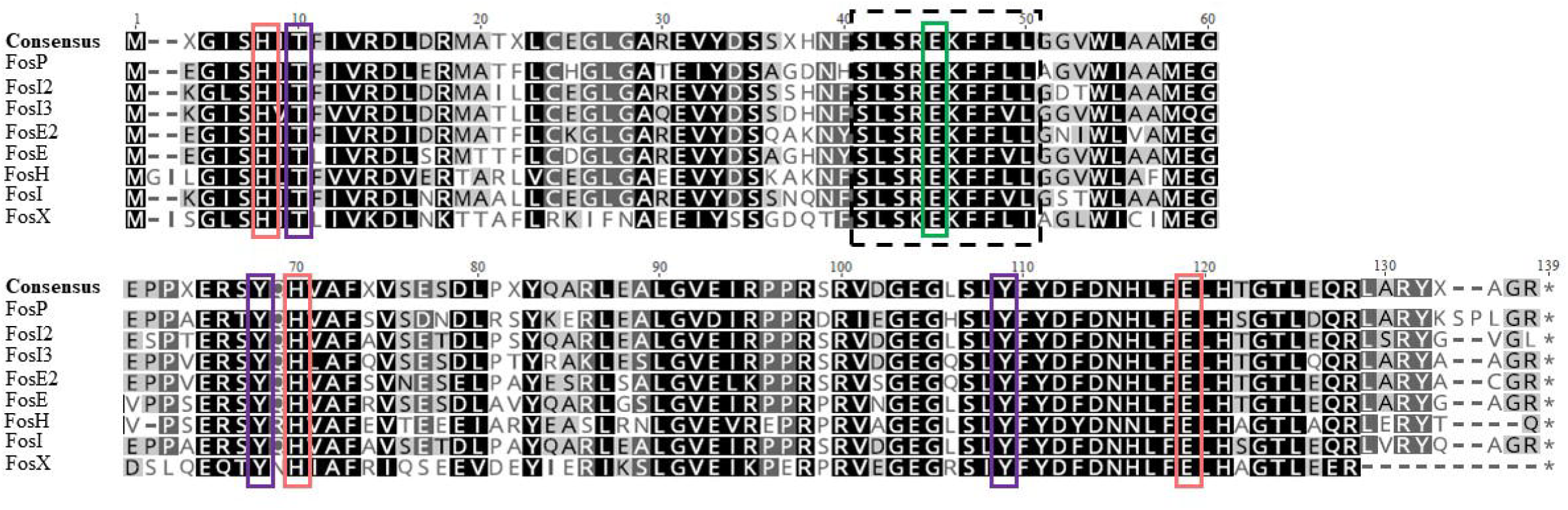
Multiple sequence alignment of newly identified fosfomycin resistance proteins compared to characterized FosX family members, including FosE, FosH, and FosI. Key functional residues are highlighted, demonstrating the conservation of crucial amino acids across the FosX family that are involved in fosfomycin hydrolysis. The dashed box highlights the conserved sequence motif (40-SLSREKFF-L/V-48), crucial for the catalytic activity of FosX-like enzymes. The green box marks the conserved E44 residue, which acts as the general base catalyst in the hydrolysis of fosfomycin. Red boxes indicate key residues involved in Mn(II) ion coordination, including histidines (H7 and H69) and glutamic acid (E118). Purple boxes highlight residues involved in hydrogen bonding, specifically tyrosine (Y108) and adjacent residues, contributing to the stabilization of the enzyme-substrate complex. The top line is the consensus sequence of all FosX family members.

### 3.2 Presence of the new *fos* genes in genomes

A close homolog to FosP (97% amino acid identity, differing by only four amino acids) was identified in an *Oxalobacteraceae* (Accession number: RYE71253) [23]. Since the accession was only 1839 nt-long, it was difficult to determine its genomic landscape. However, the presence of a different *attC* site showed that this homolog was also integron-born even if an associated integron-integrase gene could not be identified (Figure 3A). *Oxalobacteraceae* is a family within the order *Burkholderiales*, known for its diverse ecological roles and its presence in various environments, including soil and water. Several species belonging to this family are known to live in the human gut, such as *Oxalobacter* spp. [24]

**Figure 3.**
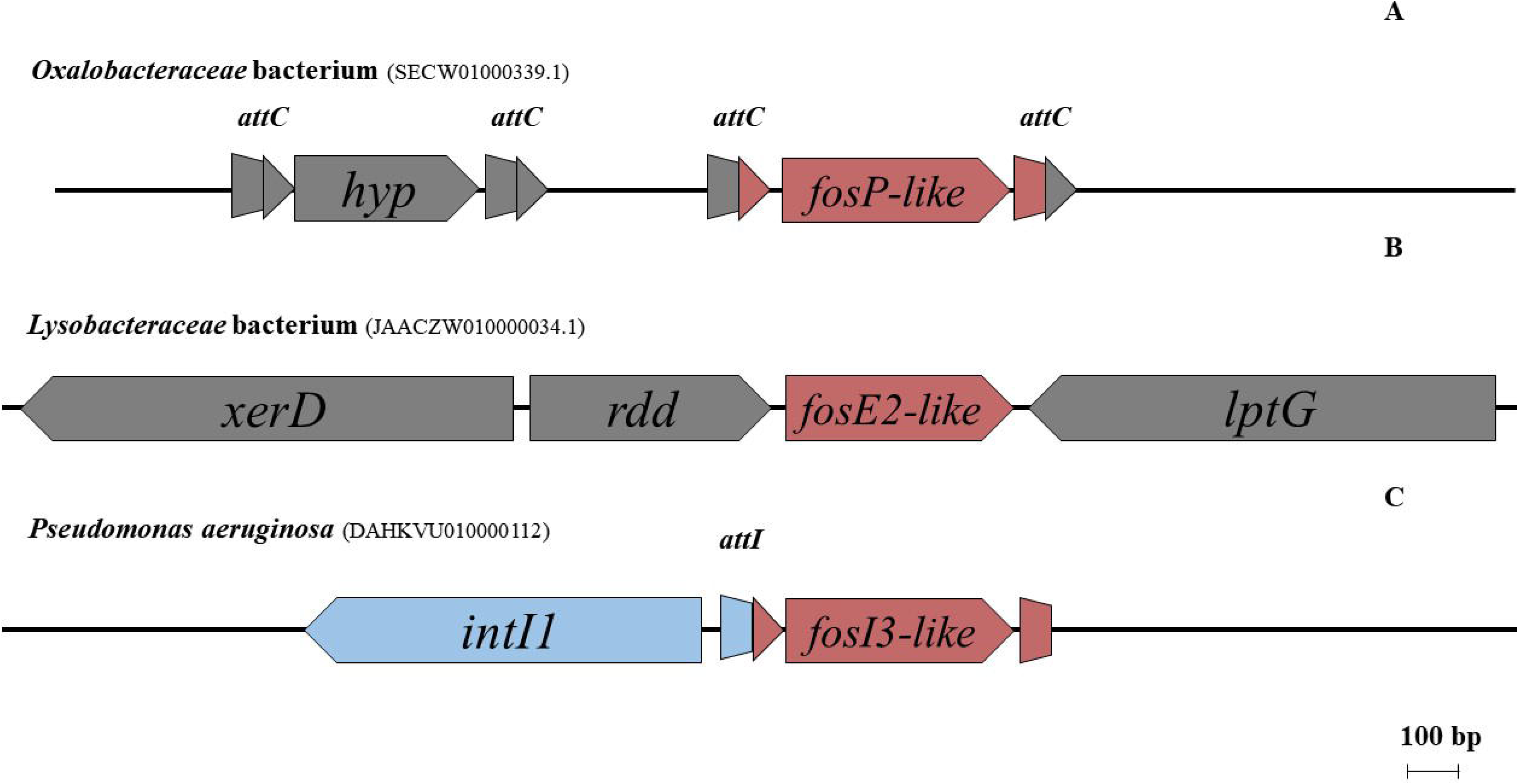
Genomic context of newly identified fos genes in various bacterial species. (A) *fosP*-like gene in *Oxalobacteraceae* bacterium flanked by *attC* sites within a class 1 integron. (B) *fosE2*-like gene in *Lysobacteraceae* bacterium located between *xerD* and *lptG*, without apparent associated mobile elements. (C) *fosI3*-like gene in *Pseudomonas aeruginosa*, positioned next to the *intI1* integrase gene within a class 1 integron.

A relatively close homolog of *fosE2* was found in the chromosome of a *Lysobacteraceae* isolate (79.9% amino acid identity). Bacteria belonging to the *Lysobacteraceae* family are known to be environmental, with some species being opportunistic pathogens such as *Xanthomonas* spp. and *Stenotrophomonas* spp. Interestingly, no *attC* site nor other mobile elements were identified in the vicinity of the *fosE2*-like gene. However, genomic analysis suggests that the *fosE* variant may have inserted itself into the genome right after the RDD family protein gene and before the *lptG* gene located downstream. This insertion could have been facilitated by the nearby *xerD* recombinase gene, although further experimentation is required to confirm its involvement. Additionally, our genomic analysis revealed a similar synteny in a *Pseudoxanthomonas* isolate, which shares the *xerD, rdd*, and *lptG* genes (with >91 % nucleotide identity) but lacks the *fosE2*-like gene (Figure S2). This led us to hypothesize that *fosE2* was inserted via an unknown mechanism, possibly involving the recombinase activity of XerD. However, further experiments are needed to confirm this hypothesis and clarify the exact mechanism of insertion.

Finally, a homolog of FosI3 (93% amino acid identity) was identified in a *Pseudomonas aeruginosa* isolate in a study presenting a new *de novo* sequence read assembler for microbial genomes [25]. Unfortunately, specific details about this isolate were not available. The *fosI3* gene was located at the first position of a class 1 integron, but the short shotgun sequence did not provide information about the subsequent gene cassettes in this array (Figure 3C).

It is of note that no close homolog of *fosI2* has been identified in the NCBI database, showing the limit of our knowledge about the current dissemination of this gene among bacterial species and the need of new sequencing data in order to have a better picture of the dissemination of this variant.

### 3.3 Functional characterization of the new Fos enzymes

The four *fos* genes were cloned into the same pMK vector, flanked by a P_*bla*_ promoter and a *rrnB* terminator, to compare their impact on fosfomycin resistance in *E. coli* Dh10β. Table 1 details the minimum inhibitory concentrations (MICs) by agar dilution and by standard disk diffusion of the pMK *E. coli* DH10β clones carrying the different *fos* genes, revealing that each of the new Fos proteins conferred clinical resistance to fosfomycin upon expression in *E. coli* ranging from 32 μg/ml to 256 μg/ml. Subsequently, the impact of phosphonoformate on inhibiting these Fos proteins was evaluated, using a FosA3-producing *E. coli* clone as a positive control. Surprisingly, 5 mM of PPF had almost no effect on the MICs for the newly identified Fos proteins, showing only a slight 2-fold decrease in most cases (and a 4-fold decrease for *fosE2*). In contrast, the positive control, *fosA3*, showed a >1000-fold decrease in MIC when exposed to PPF, demonstrating that PPF is highly effective against FosA family enzymes but not against the FosX-like proteins described in this study. Phosphonoformate, commonly known as Foscarnet, is an antiviral agent previously identified to inhibit FosA and FosB proteins effectively [22,26,27]. It acts as a transition state analogue inhibitor, binding to the Mn(II) center in FosA and likely in FosB, forming a five-coordinate complex that mimics the transition state geometry of the fosfomycin inactivation reaction. This interaction disrupts the enzyme’s ability to conjugate glutathione or bacillithiol to fosfomycin, thereby preserving the antibiotic efficacy [22,27]. The observation of a slight decrease in fosfomycin MIC in the presence of PPF for FosX-like enzymes is unexpected. FosX and FosA/B enzymes share conserved amino acid residues involved in metal ion binding and coordination, which are important for their structural stability. However, the catalytic sites of these enzymes differ significantly. In FosX-like enzymes, the key region is (40-SLSR**E**KFF-L/V-48) (Figure 2), with E44 residue playing a critical role in the hydrolysis of fosfomycin, whereas FosA and FosB rely on different catalytic residues to modify fosfomycin. One possible explanation for the slight decrease in MIC could be that PPF interacts weakly with the metal-binding residues in FosX-like enzymes. While these residues contribute to metal coordination, they are not as crucial for the hydrolytic activity of FosX enzymes as they are for FosA and FosB, where metal ion coordination is directly tied to their catalytic mechanism. In FosA/B enzymes, the metal-binding residues are essential for catalyzing the conjugation reaction that inactivates fosfomycin. In contrast, the hydrolytic activity of FosX seems to be less dependent on these metal-binding residues, though they may still enhance the enzyme’s efficiency. Therefore, the slight effect of PPF on FosX-like enzymes may be due to its weak interaction with metal-binding residues that support, but do not fully govern, the enzyme’s catalytic function. More detailed studies are necessary to clarify the underlying mechanisms of this observation.

**Table 1.**
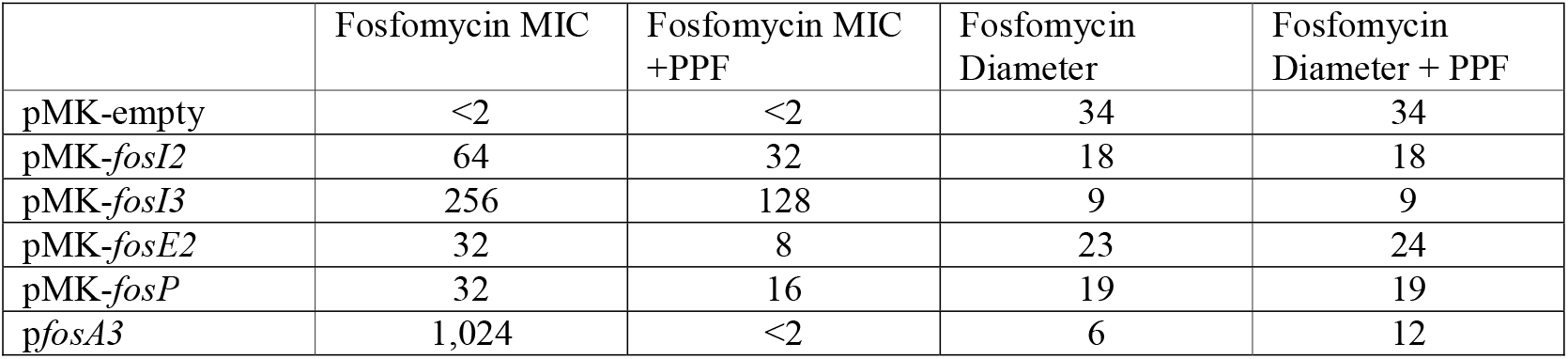
Minimum inhibitory concentrations by agar dilution and inhibition zones by disk diffusion of the pMK *E. coli* DH10β clones carrying the different *fos* genes. *pfosA3* is a pUCP24 plasmid carrying the glutathione-S-transferase gene *fosA3*, used as a positive control for the phosphonoformate (PPF) inhibition effect.

## 4. Conclusion

Our study successfully identified four novel fosfomycin resistance genes belonging to the *fosX* family in environmental samples from geographically distinct locations, ranging from highly polluted sites in India to the isolated Antarctic. While the specific variants differed across sites, *fosX*-family fosfomycin resistance genes were found in all these diverse environments. These findings indicate that acquired *fosX* resistance mechanisms are geographically widespread and are likely to confer selective advantage in various environmental contexts. It seems likely that further *fos* families and gene diversity would be revealed by additional surveys. Such studies are crucial for enhancing resistance gene databases, potentially aiding in the future development of diagnostic tools to detect *fosX*-family resistance genes in clinical settings. Indeed, one of the *fosI3* variants was found in the pathogen *Pseudomonas aeruginosa*, and raises significant clinical concerns since the presence of such resistance genes in pathogenic bacteria suggests that these mechanisms could contribute to future clinical challenges in treating infections.

In recent years, several new *fos* genes have been identified, but the majority belong to the GST or BST families and are generally associated with insertion sequences or acquired through genomic islands [28–30]. In this study, we characterized new *fos* determinants that are predominantly present within class 1 integrons. It is noteworthy that *fosX* genes are believed to originate from *Listeria* spp., a Gram-positive bacterium [3], while the acquired forms of these genes are primarily found in Gram-negative species. While transfer of resistance genes between Gram-positive and Gram-negative bacteria is known to occur, it is less common, making the presence of FosX-mediated resistance mechanisms in both groups an area worthy of further investigation.

The FosX family of enzymes remains poorly studied compared to other fosfomycin resistance mechanisms. Recent studies detected *fosX* genes in environmental metagenomes but did not characterize their impact on fosfomycin resistance or record their nucleotide sequences [31]. These enzymes warrant deeper characterization, particularly because they appear drastically less affected by phosphonoformate. This kind of resistance potentially makes FosX enzymes a greater clinical threat. While our current approach using class 1 integron gene cassette PCR amplicon was effective in identifying these genes, it has limitations in capturing the full genetic diversity and potential novel mechanisms present in bacterial genomes outside the class 1 integron genetic environment. This method amplifies specific sequences, increasing the likelihood of detecting known resistance genes but potentially missing novel elements that do not fit this profile. To address these limitations and detect a broader range of new resistance determinants, future studies could focus on performing functional metagenomics from the whole genomes of resistant bacteria that are selected on specific media. This approach narrows the focus to bacteria that can grow under the selective conditions, ensuring that resistance genes of interest are enriched within the bacterial population. While this method excludes bacteria that do not grow in these settings, it allows for more targeted analysis of the genomes of resistant strains[32].

## Competing Interests

The authors do not have any conflict of interest.

## Funding

This work was funded by; the Swedish Research Council (VR), Contract No. 2018-05771 and 2022-00945, and the UGOT Challenges Initiative at the University of Gothenburg

## Ethical approval

Not required

## Sequence information

Gene sequences were lodged under the following accession numbers: PQ186074 (*fosE2*), PQ186075 (*fosI2*), PQ186076 (*fosI3*) and PQ186077 (*fosP*)

## Notes

### Competing Interest Statement

The authors have declared no competing interest.

