## Supplemental Figure 1 for "Identification of Novel FosX Family Determinants from Diverse Environmental Samples"

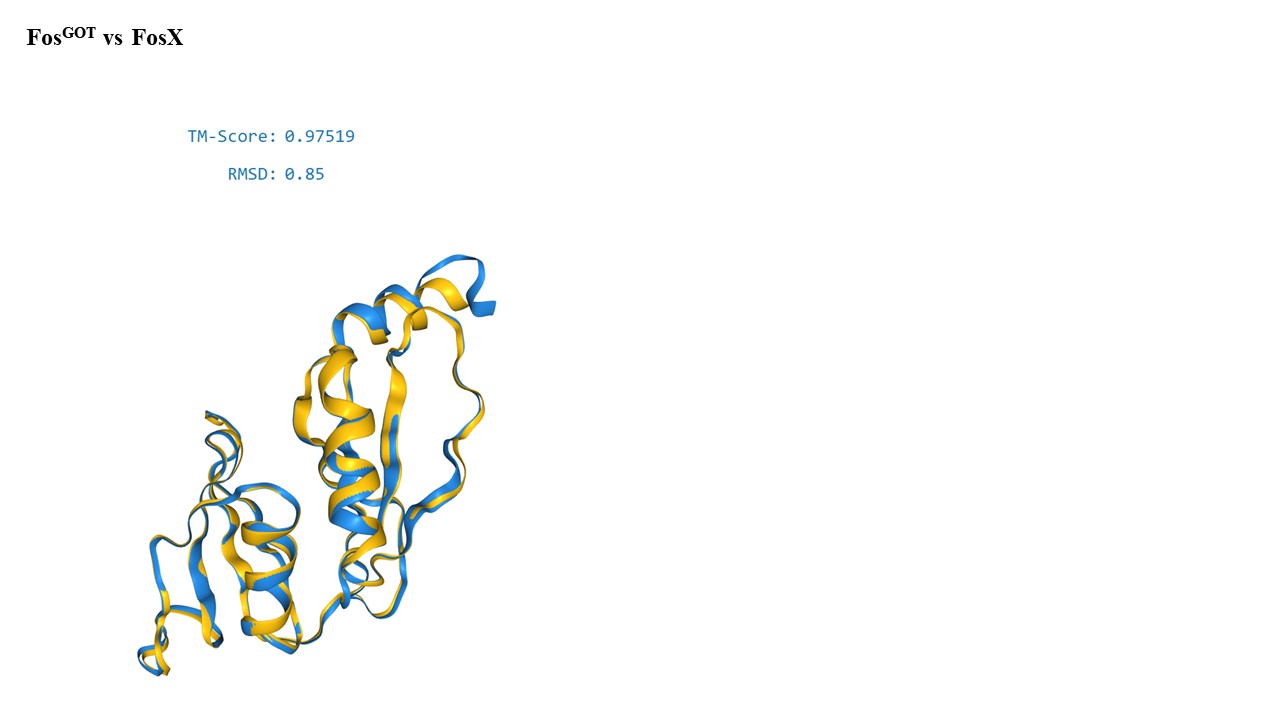

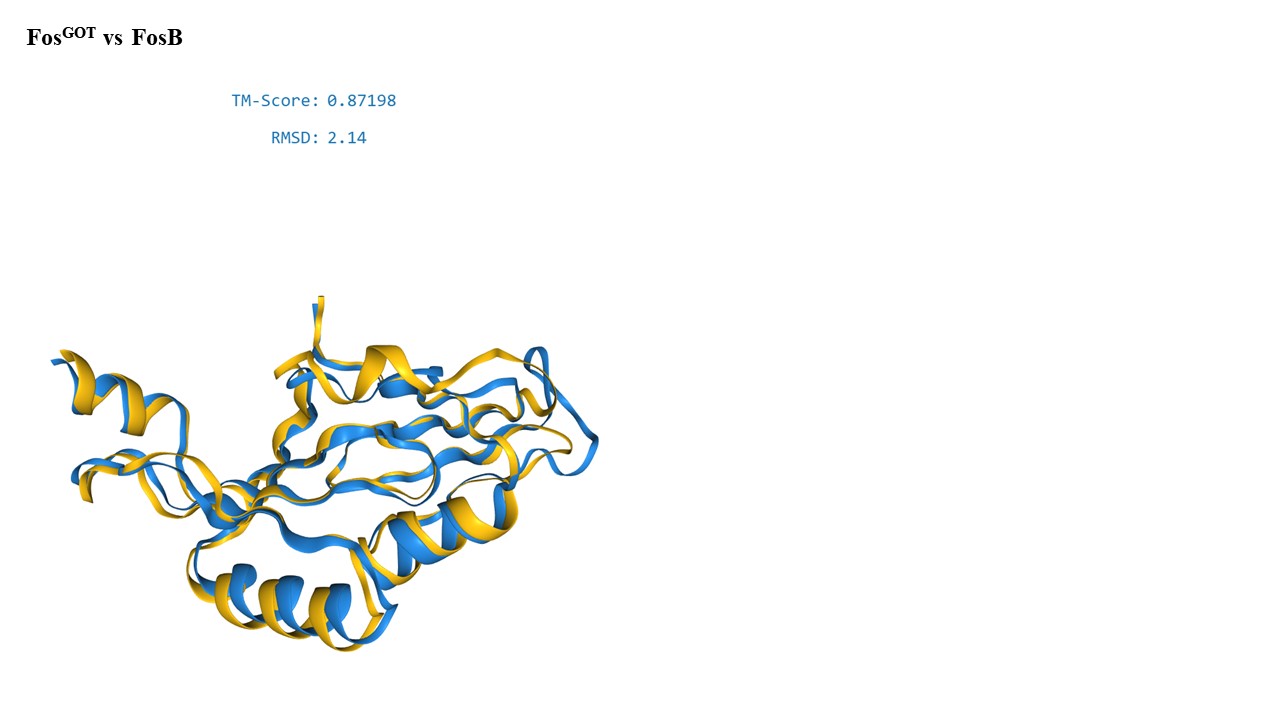

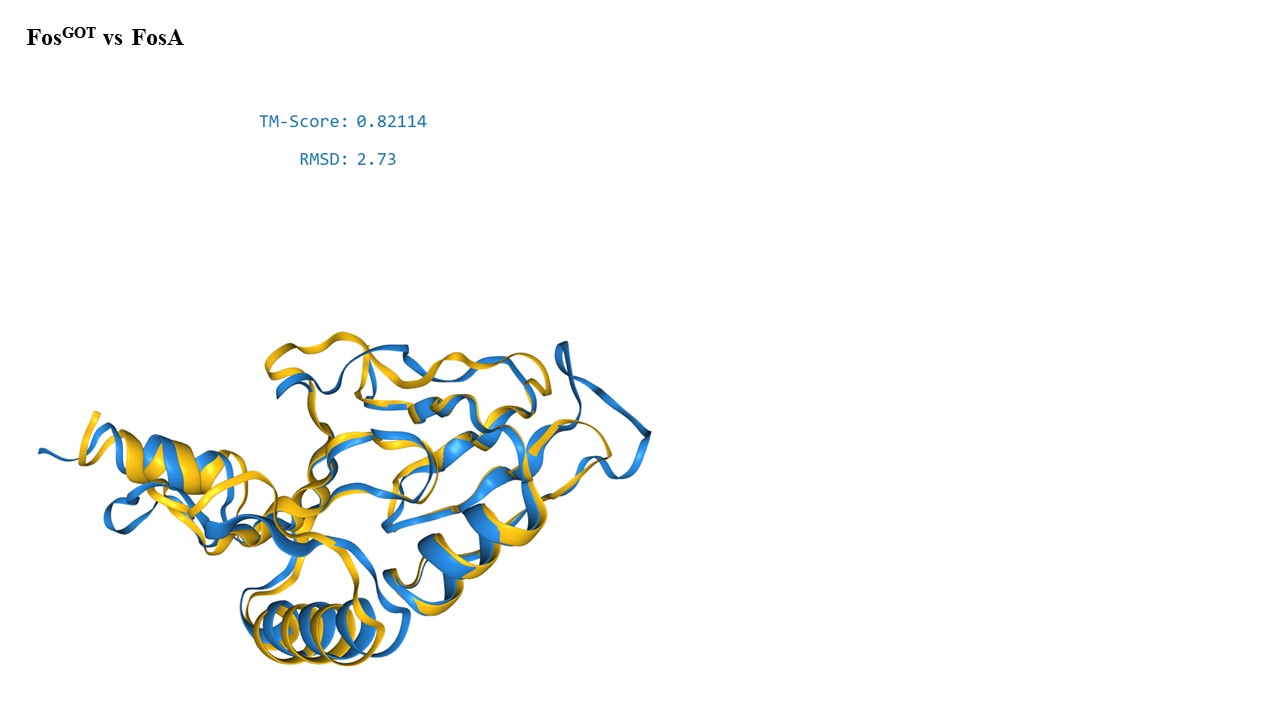

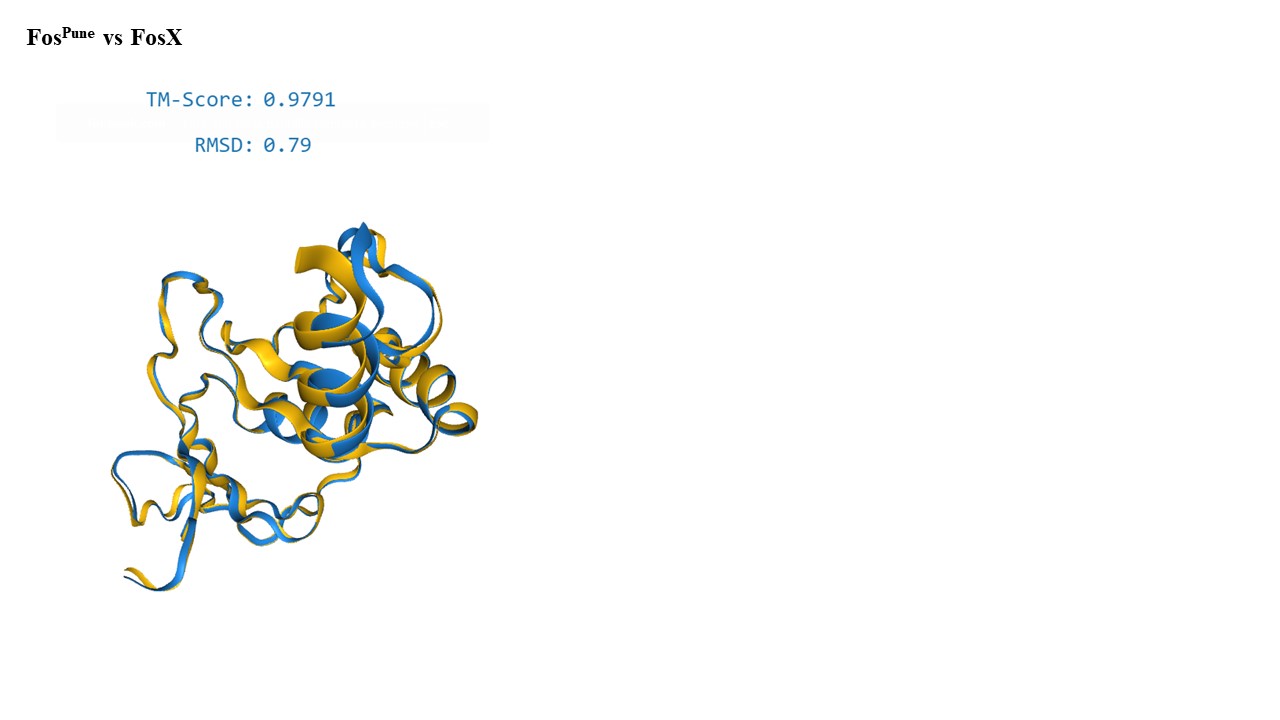

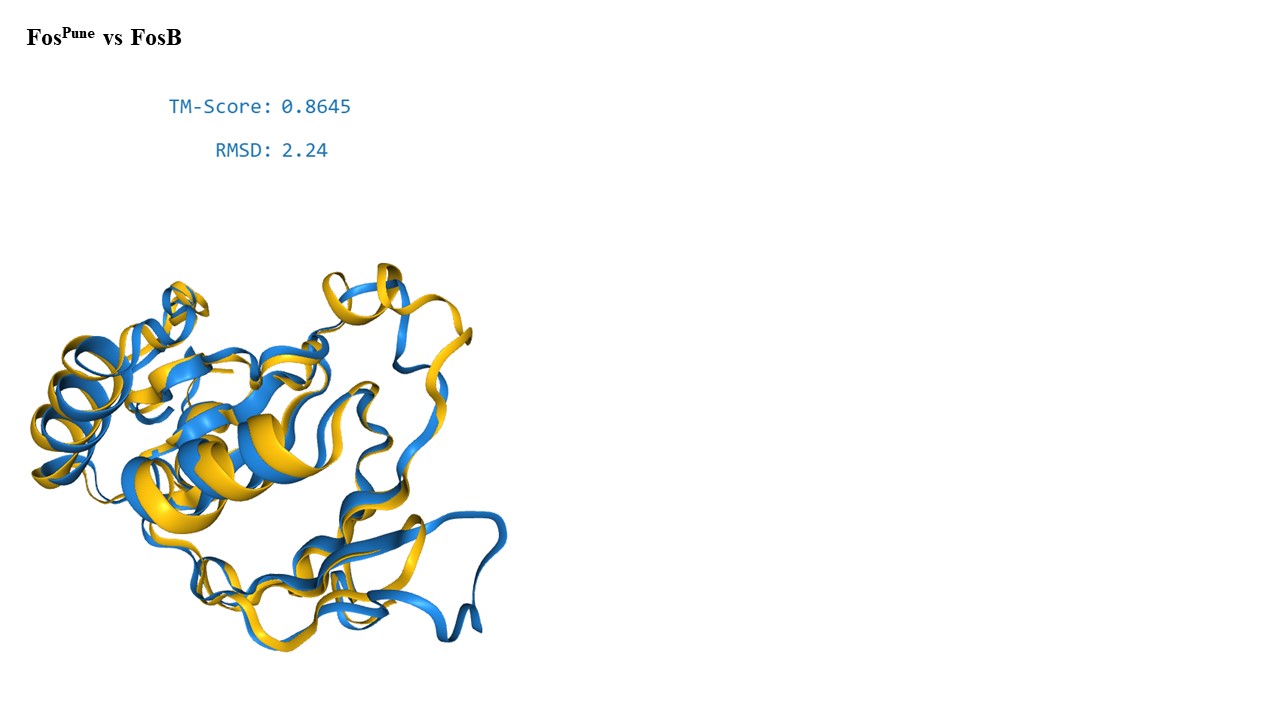

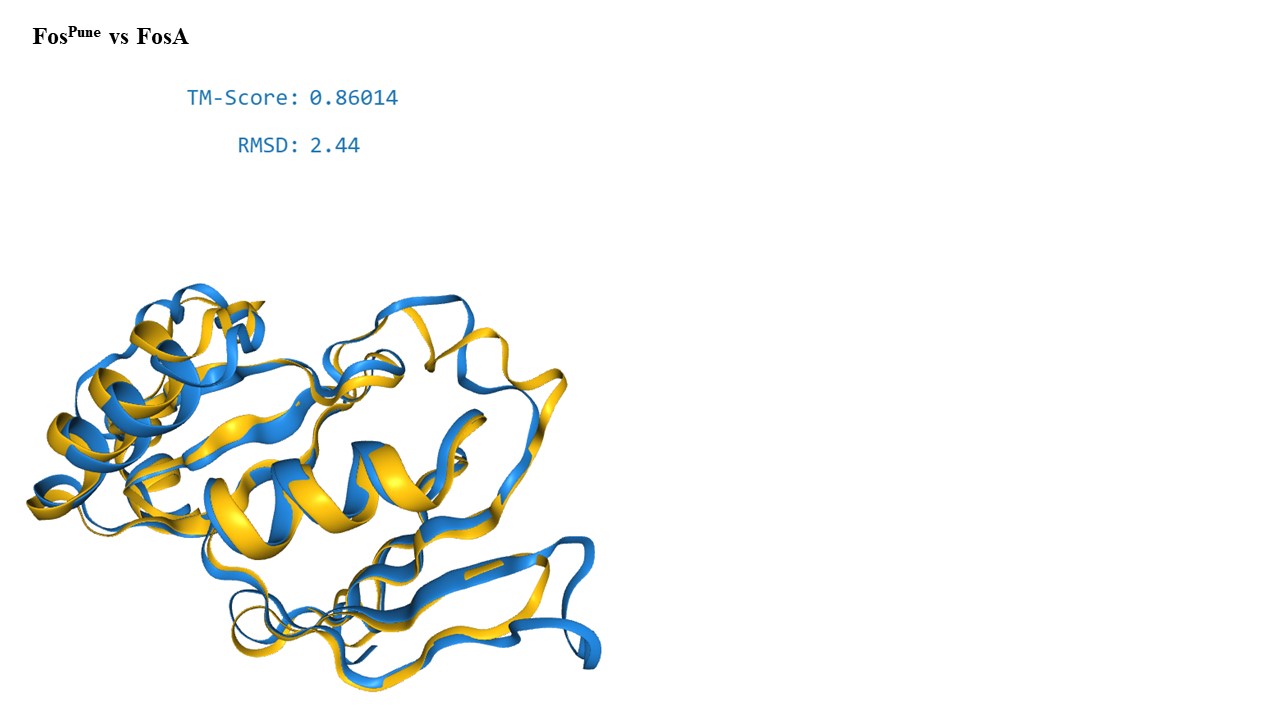

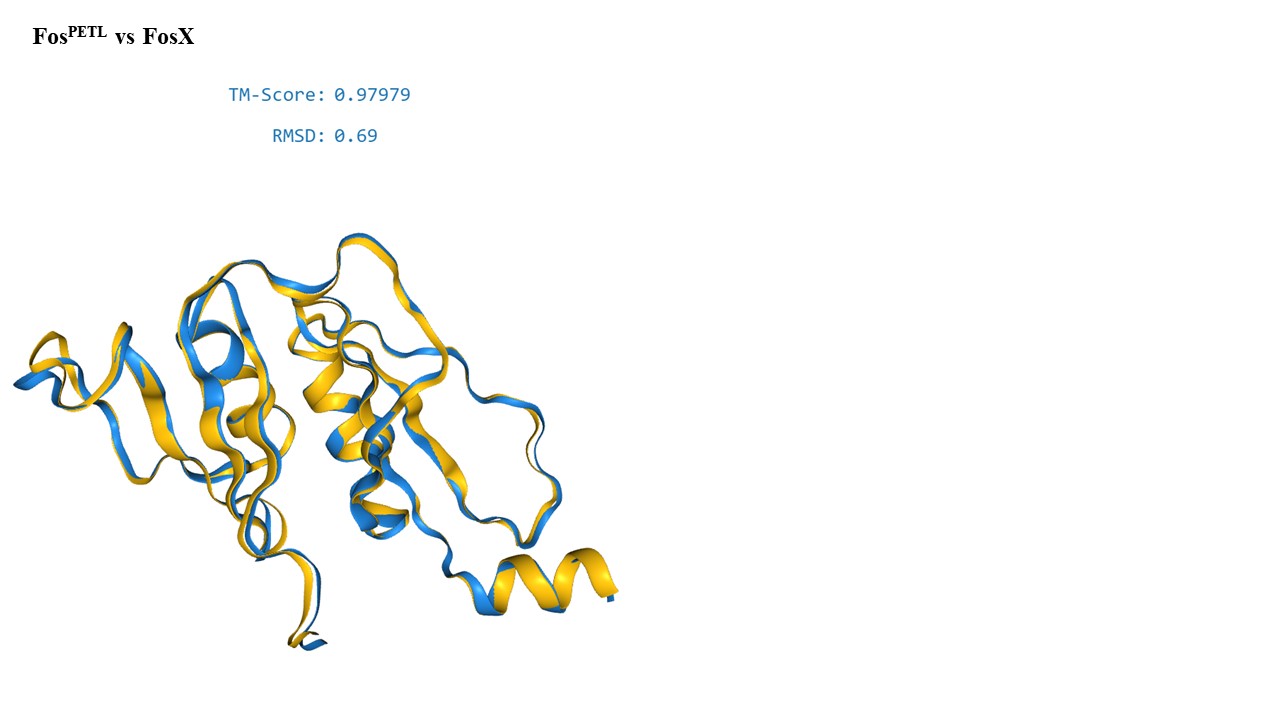

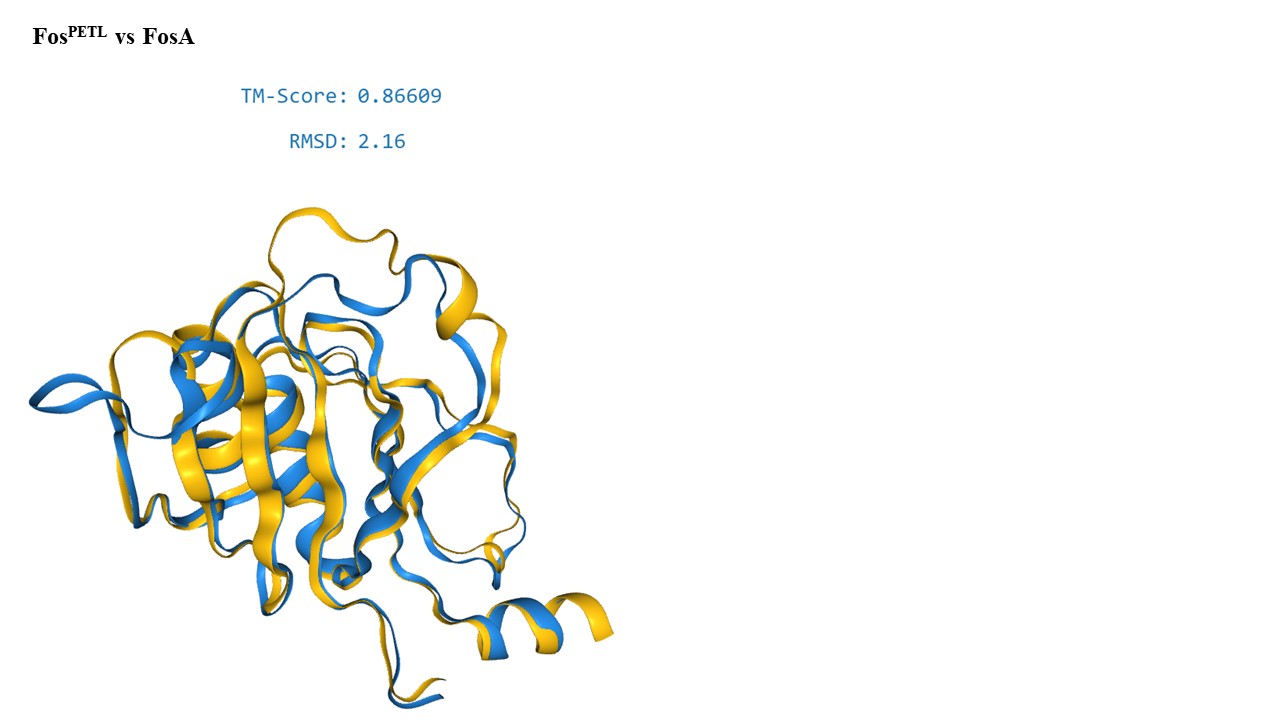

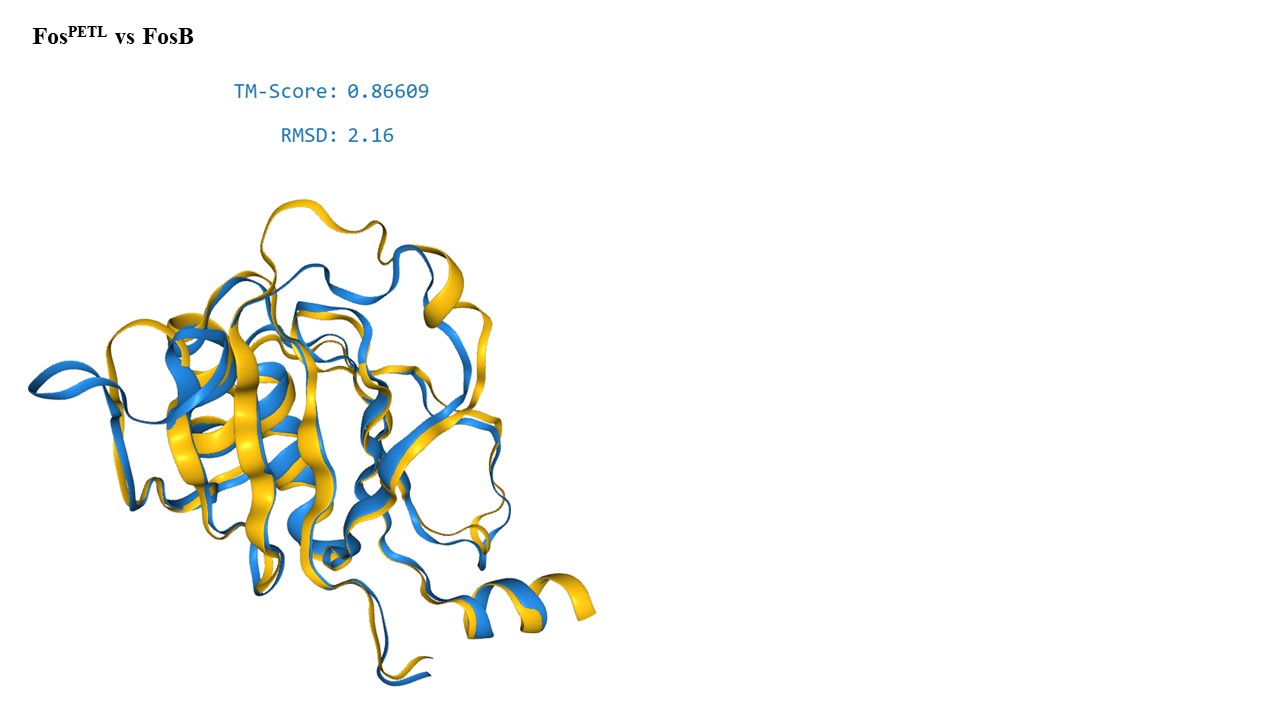

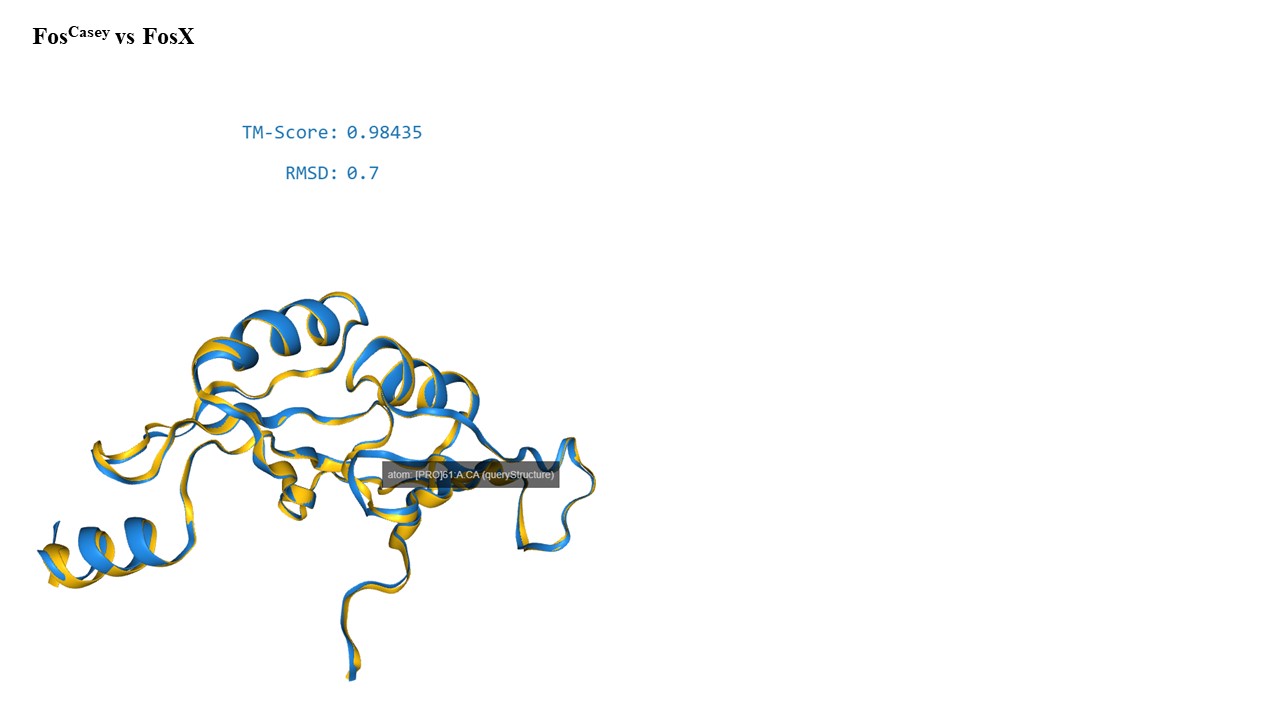

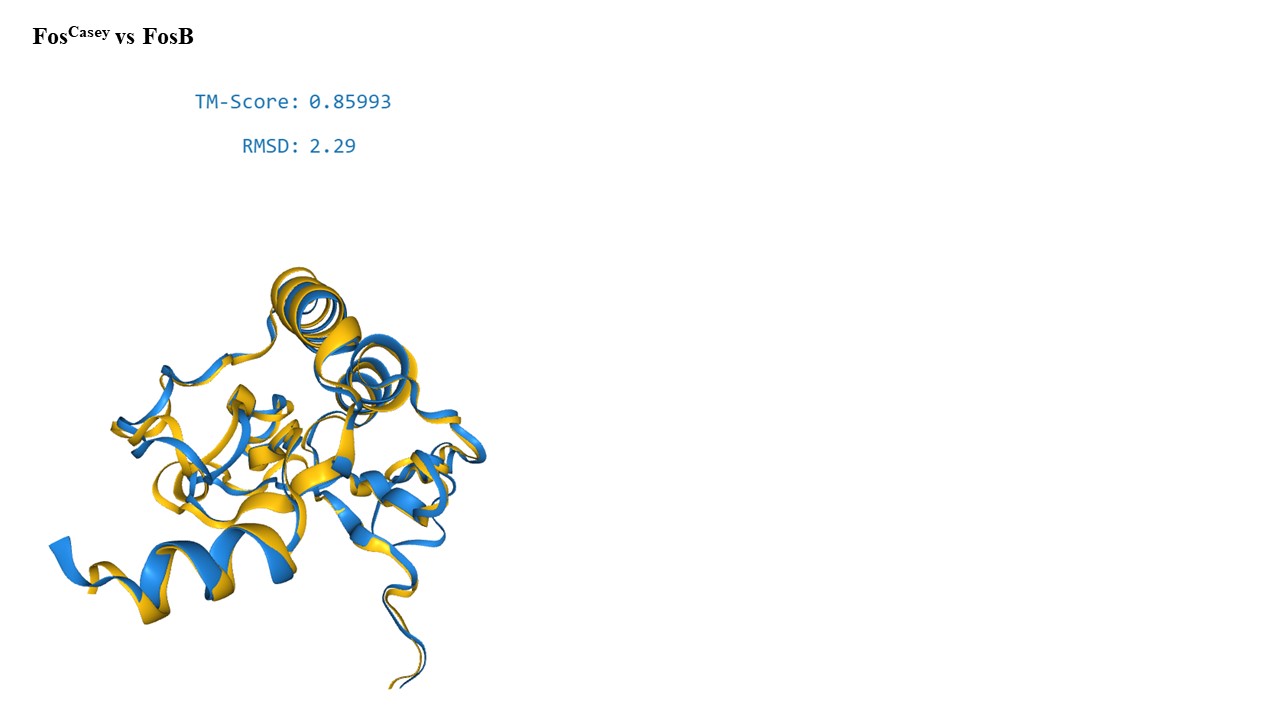

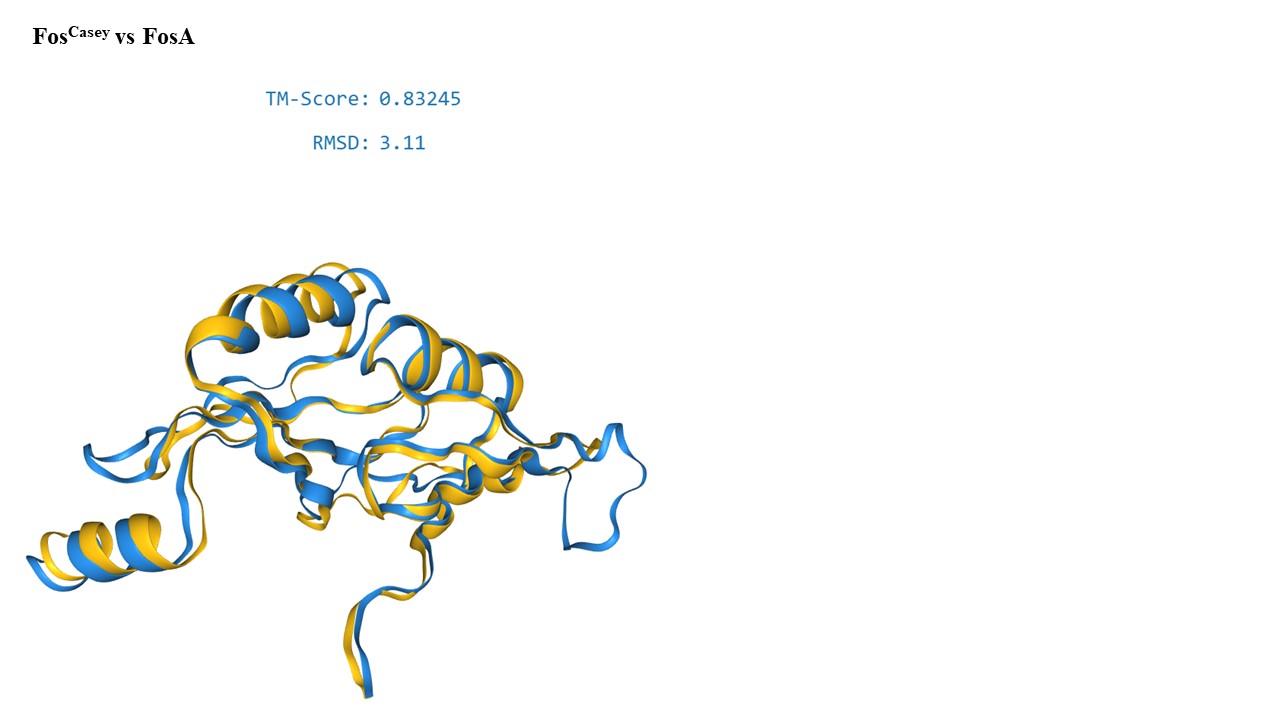

**Figure S1**. Three-dimensional structure alignment using AlphaFold2. TM-score and RMSD values are used to assess the structural similarity between the newly identified Fos proteins and known FosA, FosB, and FosX enzymes Structural alignment of the newly identified Fos proteins with FosA, FosB, and FosX enzymes. The TM-scores and RMSD values indicate the degree of structural similarity, with TM-scores above 0.5 and RMSD values below 2 Å suggesting a significant structural match. The high TM-scores (>0.97) and low RMSD values (<0.8) between the new Fos proteins and FosX enzymes confirm their close structural relationship, supporting their classification within the FosX family. Accession numbers: FosA (Q9I4K6), FosB (Q2FVT3) and FosX (Q8Y6I2).
