## Supplemental Figure 2 for "Identification of Novel FosX Family Determinants from Diverse Environmental Samples"

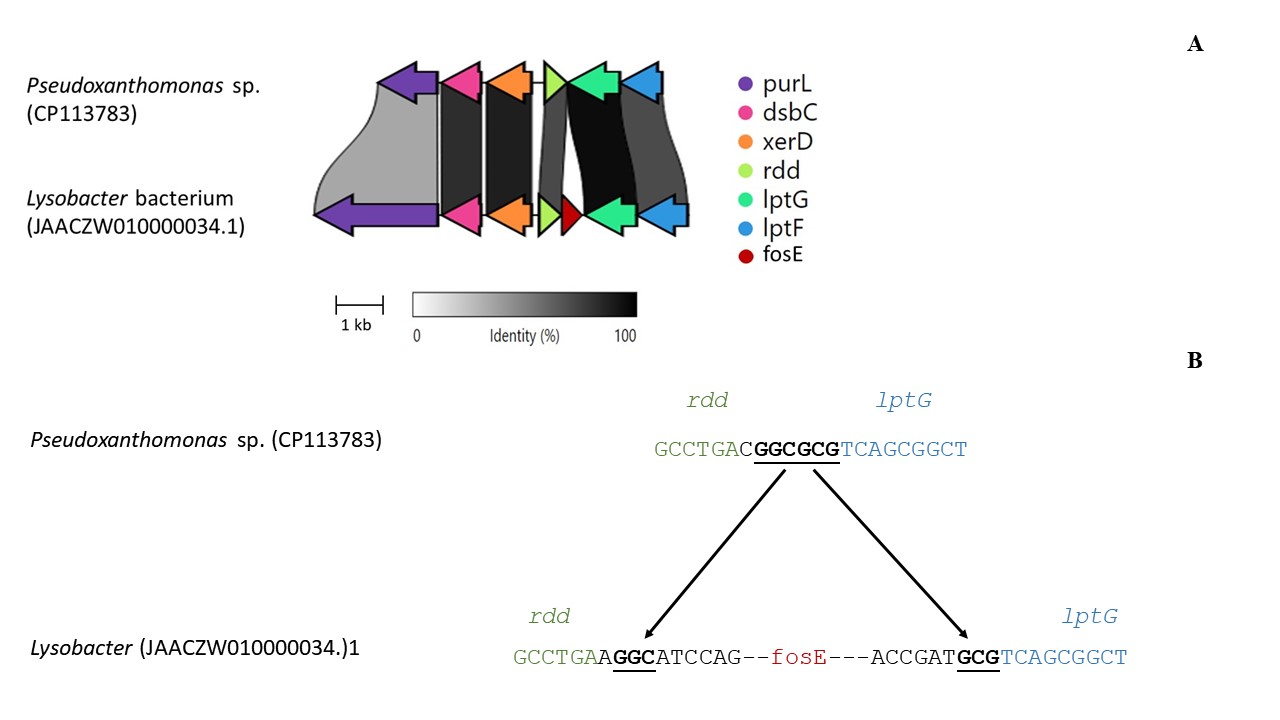

**Figure S2.** (A) Synteny comparison between Pseudoxanthomonas sp. (CP113783) and Lysobacter bacterium (JAACZW010000034.1), showing the conserved gene arrangement in the vicinity of the ***fosE*** gene. The grayscale shading represents nucleotide identity between the two sequences. (B) Detailed alignment of the genomic sequences around the ***rdd*** and ***lptG*** genes in Pseudoxanthomonas and Lysobacter, highlighting the insertion site of the ***fosE*** gene in Lysobacter (red). The ***rdd*** gene is labeled in green, and the ***lptG*** gene in blue, emphasizing the conserved regions in both organisms. The underlined sequence represents the potential insertion site of the *fosE* region.
